# Top-down influences on neural music processing: preference, enjoyment and familiarity influence neural tracking and brain rhythms differently

**DOI:** 10.64898/2026.07.10.737714

**Authors:** Saara M. Varjopuro, Rosanne H. Timmerman, Tanja Atanasova, Sarah C. Allen, Stratos Koukouvinis, Anne Keitel

**Author notes:** Corresponding authors: Anne Keitel and Saara M. Varjopuro. Shared first authors (Saara M. Varjopuro, Rosanne H. Timmerman, Tanja Atanasova).

## Abstract

Music enjoyment and familiarity are closely related but often confounded in studies of neural music processing. Here, we investigated their distinct contributions to cortical oscillatory activity and neural tracking of music using electroencephalography (EEG). Thirty-two participants listened to self-selected all-time favourite songs, recent favourite songs, and tempo-matched songs from disliked genres. This novel paradigm dissociated familiarity from enjoyment by including highly enjoyed songs that differed in familiarity. Spectral power and cortical tracking (using Mutual Information) were analysed using linear mixed-effects models with enjoyment and familiarity ratings. Familiarity was associated with increased left-frontal alpha power, whereas enjoyment predicted increased theta and beta power, demonstrating distinct oscillatory signatures for these dimensions. An interaction revealed that the positive relationship between enjoyment and theta power was strongest for highly familiar music. Cortical tracking analyses showed that greater enjoyment was associated with reduced delta-band tracking, with a significant interaction indicating that this negative relationship was present for highly familiar songs but not for less familiar songs. These findings indicate that enjoyment and familiarity differentially shape neural responses to music and highlight the importance of modelling both factors to disentangle their distinct effects on neural activity during music listening.

## Introduction

Music has been part of human life all over the world for thousands of years. It surrounds us daily and listening to music can have stress-reducing effects (Linnemann et al., 2016), reduce pain (Costa et al., 2018; Jafari et al., 2012), and decrease depression and anxiety (Costa et al., 2018; Leubner & Hinterberger, 2017; Särkämö et al., 2008). Moreover, daily music listening can induce greater grey matter reorganisation (Särkämö et al., 2014) which highlights the potentially profound effect of music on the brain.

Music preference is influenced by two broad factors: music-internal and music-external parameters (Schäfer & Sedlmeier, 2010). Music-internal factors include elements such as tempo, harmony, and complexity, while music-external factors relate to the listener’s characteristics, including age, gender, personality, and cultural background. These preferences are highly personal (Huang et al., 2020; McDermott et al., 2014; Schaefer, 2016). A study with infants indicated a preference for music that is common in the culture (Soley & Hannon, 2010). Familiar music is also often preferred, sometimes referred to as a ‘mere exposure effect’ (Peretz et al., 1998; Madison & Schiölde, 2017). Music appreciation strongly relies on memory processes and familiarity and has therefore been described as an implicit memory phenomenon (Peretz et al., 1998; Green et al., 2012). Another explanation for our preference for familiar music comes from predictive processing, a theory according to which the brain creates a model to predict incoming sensory input (Fitzgerald & Todd, 2020), and its connections to the reward system. It has been shown that the better the sensory input matches the prediction, the stronger the response from dopamine cells, resulting in more pleasure (Salimpoor et al., 2015). Thus, more familiarity with a musical piece would involve more accurate predictions, resulting in a more pleasurable experience.

Cortical oscillatory activity can shed light on how the brain processes music, and effects of preference, enjoyment or familiarity on music processing have been shown in all traditional frequency bands (i.e., delta (0–4 Hz), theta (4–8 Hz), alpha (8–14 Hz), beta (14–30 Hz), and gamma (>30 Hz)). Frontal theta power has been found to increase when listening to pleasant music (Ara & Marco-Pallarés, 2020; Sammler et al., 2007) and this has also been found to be stronger when listening to a favourite song compared to a moderately preferred song (Tseng, 2021). Another study found that posterior alpha power increased when listening to a preferred genre compared to a nonpreferred genre (Hurless et al., 2013). When listening to self-selected preferred music, an increase in posterior alpha and beta as well as temporal gamma power has been found (Nakano et al., 2021). These studies tend to convolute enjoyment and familiarity when self-selected preferred music, which is both enjoyed and familiar, is used to study enjoyment effects.

When the specific effect of music familiarity on oscillatory power was studied, one study found differences in all frequency bands (Thammasan et al., 2016). Specifically, listening to familiar music was associated with stronger frontal delta and posterior alpha power, as well as lower frontal midline/temporal theta, and lower temporo-occipital beta and gamma power, than when listening to unfamiliar music. Another study found different connectivity patterns in the theta band for pleasant familiar and pleasant unfamiliar songs (Ara & Marco-Pallares, 2021). Overall, these varied results indicate that preference, enjoyment and familiarity of music might have distinct effects on cortical music processing.

Beyond brain oscillations, cortical activity has also been found to track slow amplitude fluctuations in the acoustic signal such as rhythm and tempo (Keitel et al., 2025; Osorio & Assaneo, 2025). Cortical tracking of music might also be influenced by enjoyment and familiarity, although previous findings are mixed. Enjoyment has been associated with stronger cortical tracking in one study (Keitel et al., 2025), but not in another (Weineck et al, 2022). Similarly, familiarity with music has been associated with stronger neural tracking in some studies (Doelling & Poeppel, 2015; Weineck et al., 2022), and less cortical tracking in others (Kumagai et al., 2017, 2018). Similar to the studies examining oscillatory power discussed above, research on enjoyment and familiarity has typically not been designed in a way that allows their effects to be disentangled.

Previous studies looking at preferred compared to nonpreferred music have either used pre-selected songs (Ara & Marco-Pallarés, 2020; Ara & Marco-Pallarés, 2021; Sammler et al., 2007; Hurless et al., 2013) or have asked participants to bring their own music, with the nonpreferred songs being sourced from other participants’ music (Nakano et al., 2021). Similarly, studies looking at familiarity have not always considered music preference (Thammasan et al., 2016). The current study dissociated between familiarity and enjoyment by presenting participants with their “all-time favourite” music, which listeners were very familiar with, and their “recent favourite” music, which listeners were less familiar with, while showing comparable enjoyment. These preferred stimuli were tempo-matched with songs from a disliked genre, thus tailoring the stimuli to each participant’s individual preferences. We analysed oscillatory activity and cortical tracking comprehensively across a wide range of frequencies and tested the hypotheses that listening to favourite songs would result in higher power (in theta, alpha, beta and gamma bands) compared to disliked songs. The novel paradigm used in this pre-registered study (https://osf.io/9yuhx) allowed us to better disentangle familiarity and enjoyment effects on spectral activity and cortical tracking processes during listening to preferred and unpreferred music.

## Methods

### Participants

Thirty-two participants took part in the study (17 female; age range 19–27 years, *M* = 22.2, *SD* = 2.5). Participants were native English speakers, with no previous diagnosis of psychological or neurological disorders. All participants were right-handed (Oldfield, 1971). Thirty-one participants reported normal hearing (Quick Hearing Check; Koike et al., 1994), while one reported a score that might suggest slightly diminished hearing (score of 23/60, hearing test recommended from score 20). Participants also completed a short version of the Goldsmith’s Musical Sophistication Index (Gold-MSI; Müllensiefen et al., 2014). On average, participants scored a 4.26 (out of 7) on this questionnaire (range = 2.27–6, *SD* = 1.11). Seven participants reported no musical training, while six participants reported 10 or more years of musical training.

This study was pre-registered on the Open Science Framework in April 2024 (https://osf.io/9yuhx). Any deviations from the pre-registered analyses are detailed where appropriate. This study was approved by the School of Social Sciences Research Ethics Committee at the University of Dundee (ethics UoD-SoSS-PSY-UG-2021-265). The research was conducted in accordance with the Declaration of Helsinki, and all methods were performed in accordance with the relevant guidelines and regulations. Participants gave written informed consent prior to participation and were given a £20 shopping voucher for participating.

### Music stimuli

Participants were asked to provide their favourite songs and disliked genres via an online session before coming into the lab. We requested them to submit two ‘all-time’ favourite songs and two ‘recent’ favourite songs. Additionally, we requested participants to choose up to three disliked genres from the following list: blues, classical, country, electronic, folk, funk, heavy metal, hip hop, jazz, latin/reggaeton, new age, pop, punk, R&B, rap, reggae, rock, soul or world. Four songs were chosen as the musical stimuli: 1) an all-time favourite song, 2) a recent favourite song, 3) a song from the disliked genres that matched the tempo and peak in the modulation spectrum of song 1, and 4) a song from the disliked genres that matched the tempo and peak in the modulation spectrum of song 2. To choose disliked songs, we first obtained the songs’ BPMs online (www.songbpm.com and www.getsongbpm). This allowed us to identify songs that likely matched the tempo from the favourite songs. Once a potential match was found, the pair was compared using the temporal modulation spectrum (Ding et al., 2017). The highest accepted difference in the songs’ peak frequencies was 1 Hz (please note that the pre-registration specifies a maximum difference of .1 Hz, which was a typo). All songs had a sampling rate of 44,100 Hz and were cut to two minutes in length. A list of all songs is available in the Supplementary Information (supplemental Table S1).

### Experimental procedure

The experiment consisted of two sessions: an online and an in-person session. The online experiment was created in Gorilla Experiment Builder (Anwyl-Irvine et al., 2020), to collect participants’ favourite songs and disliked genres. It also contained screening questions to assess whether the participant met inclusion criteria. We collected information on age, sex, handedness, neurological and psychological health. During the in-person session, participants completed the hearing questionnaire (Quick Hearing Check, Koike et al., 1994) and provided information about the amount of sleep and alcohol consumption the night before testing, caffeine intake the day of testing, plus their average sleep hours and average caffeine consumption. This information was used to confirm that participants were fit for testing.

The EEG recording took place in a soundproof booth. Participants were seated at approximately 65 cm from a 24” computer screen (‘iiyama ProLite B2483HS’). Music was played through high-quality, wired headphones (‘Sennheiser HD25 700’). Auditory and visual information were presented using Psychtoolbox Version 3.0.17 for MATLAB (Brainard, 1997). Instructions with white text on black background were given on-screen. Participants were instructed to sit as still as possible and look at a green circle in the middle of the screen while listening to the songs. The order of the songs was randomised for each participant. Self-reported enjoyment and familiarity ratings were collected after listening to each of the four music tracks. Enjoyment and familiarity ratings were measured using Visual Analog Scales (VAS): a 10 cm horizontal line on paper presented with ’did not enjoy at all’ and ’enjoyed a lot’ as the two ends on the spectrum for the enjoyment question, and ’not at all familiar’ and ’extremely familiar’ for the familiarity question. Participants were instructed to draw a short vertical line on the scale, depending on their subjective experience. The lines were scored from 0–100.

### EEG data acquisition and preprocessing

EEG recordings were obtained with a 64-channel BioSemi ActiveTwo system at a sampling rate of 512 Hz. The electrodes were placed according to the International 10-20 system. Eye-movements were recorded throughout the experiment using four electrooculographic (EOG) electrodes placed at the outer canthus of each eye (for horizontal movements) and above and below the left eye (for vertical eye movements and blinks). Any electrodes with offsets outside ±20 mV were adjusted before the start of data recording.

Preprocessing of the EEG data was done in MATLAB R2023b using Fieldtrip (Oostenveld et al., 2011) and custom-built functions. Data was cut out for each song with a leading and trailing window of ±2 s (resulting in segments with a length of 124 s per song). After re-referencing to channel Cz, data were filtered using a high pass filter of 0.1 Hz and a low pass filter of 80 Hz (3^rd^ order Butterworth filter, forward and reverse). To identify noisy channels, we used a custom-built function that assesses the standard deviation of EEG timeseries and highlights channels that exceed thresholds across channels and within-channels over time. After visually confirming exclusions, an average of *M* = 6.41 (*SD* = 3.09) channels were interpolated (with no differences across conditions, all *p*_FDR_-values > 0.40). After re-referencing to the channel average (Bertrand et al. 1985), an Independent Component Analysis (ICA) was carried out on the combined data from all conditions to identify eye movements and blinks. On average, *M* = 1.78 components (*SD* = 0.83) were removed per participant. Furthermore, a 50-Hz discrete Fourier transform filter was applied to remove line noise. For frequency analyses, noisy segments were also identified and removed by segmenting the data across all conditions into 3-s nonoverlapping windows and by removing segments with a z-scored standard deviation higher than two. An average of *M* = 6.5 (*SD* = 2.38) segments were removed across channels and participants.

### Frequency analysis

Spectral power was calculated using Fast Fourier Transform (FFT) on the segmented and pre-processed data. Specifically, we separately calculated the power for frequencies between 0.5 and 60 Hz (in semi-logarithmic frequency steps; 0.5–15 Hz: 0.25– Hz-steps, 15–30 Hz: 0.5– Hz-steps, 30–60 Hz: 1-Hz steps) using a DPSS multitaper (with ±2 Hz spectral smoothing and zero-padding to 4-s length), for every participant and condition. Power within each participant was normalised by subtracting the average power across all conditions from the power of each individual condition (mean centering).

### Music envelope preprocessing

The amplitude envelope was extracted from all music pieces using an established procedure (e.g., Keitel et al., 2018; Gross et al., 2013). Specifically, audio signals were first filtered into eight frequency bands, ranging from 100 to 8000 Hz, using a third-order Butterworth filter applied in both forward and reverse directions. These frequency bands were spaced according to equal intervals along the cochlear frequency map (Smith & Delgutte, 2007). The wideband music envelopes were obtained by averaging the magnitude of the Hilbert transformed signals across the eight bands and both audio channels. Envelopes were then down-sampled to a sampling rate of 150 Hz.

### Mutual information analysis

We calculated Gaussian Copula Mutual Information (MI; Ince et al., 2017) between the continuous EEG signals (down-sampled to 150 Hz) and the music envelopes and their derivative. This was done in the frequency domain, using a continuous wavelet transform for logarithmically-spaced frequencies between .25 and 20 Hz (Chalas et al., 2022). All signals were L1 normalised (Chalas et al., 2022). MI was initially computed at 12 different music-brain lags (from 8 to 30 sampling points, in 2-sampling point steps, corresponding to a time window between ∼50 ms and 200 ms after music onset) for each participant and condition. The optimal lag per participant and condition was chosen based on the highest MI in the delta band (i.e., MI-values averaged between 0.5 to 4 Hz across channels), as acoustic modulations in music are most prominent and relevant in the delta band (Ding et al., 2017). Only MI computed at the optimal lag for each participant and condition was used for further analyses. We chose mutual information because it is a powerful framework for cortical tracking analysis. MI quantifies stimulus-brain dependencies without assuming a linear mapping, allowing it to detect nonlinear relationships that conventional approaches may systematically overlook. Additionally using a linear analysis approach, such as temporal response functions (TRFs), is beyond the scope of this study.

### Statistical analyses

#### Outlier treatment

Behavioural and neural outlier values were removed before doing statistical analyses. For behavioural ratings, values larger/smaller than ± 3 SDs from the mean within each music condition were excluded (no exclusions for enjoyment ratings, 2 exclusions for familiarity ratings). Likewise, for neural power and MI values, outliers exceeding ± 3 SDs from the mean were removed. On average, *M* = 0.86 (*SD* = 0.51) power values were removed across channels, frequencies and conditions. For MI values, an average of *M* = 0.21 (*SD* = 0.42) values were removed across channels and conditions.

#### Comparison of behavioural ratings

The ratings for familiarity and enjoyment were compared across all conditions using two-sided dependent-samples *t*-tests. All *p*-values were corrected for multiple comparisons using FDR correction (within the set of familiarity comparisons, and within the set of enjoyment comparisons). Additionally, we analysed the relationship between enjoyment and familiarity using a linear mixed-effects model. Specifically, the model formula was *enjoyment rating ∼ familiarity rating + (1 | Participant ID)*. Note that the preregistration specified to analyse the “correlation between familiarity and enjoyment”, but that a linear mixed-effects model was deemed more in line with the overall analytical approach of the study.

#### Comparisons between music conditions

Spectral power and MI were first compared across the four conditions using two-sided dependent-samples *t*-tests. For spectral power comparisons, cluster-based permutation tests were performed to identify spatial (across all electrodes) and spectral (across frequencies between 0.5 to 60 Hz) clusters. We used a Monte-Carlo permutation approach with 5,000 randomisations and an alpha threshold of *p* = .05 (both for entering a sample into a cluster and as permutation-based significance threshold). The minimum number of adjacent electrodes was three, and of adjacent frequency bins was five (corresponding to a 1-Hz window in low frequencies, and a 4-Hz window at higher frequencies, due to the semi-logarithmic frequency resolution).

For MI, we averaged values into delta (0.25–4 Hz) and theta bands (4–8 Hz). This was done because the tempo of songs varied widely within and across participants (see Supplementary Table S1). As cortical tracking of music strongly follows the rhythmic properties of the stimulus, including tempo, a frequency-wise comparison of stimuli with different tempi seems implausible. MI was therefore compared across conditions for averaged delta and theta bands. This means the cluster-based permutation correction was only used to identify spatial clusters. All other parameters were identical to the ones used for comparison of spectral power (i.e., Monte-Carlo approach with 5,000 randomisations, alpha thresholds of *p* = .05., minimum of three electrodes). To provide an effect size for statistical tests, we report Cohen’s d for the peak electrode/frequency.

#### Linear mixed-effects models with participant ratings

To find out whether individual participant ratings predicted spectral power and MI, we used linear mixed-effects models. For spectral power, a linear mixed-effects model was fitted for each channel and frequency, with (*z*-scored) power as dependent variable. Included fixed effects were the *z*-scored continuous ratings for enjoyment and familiarity and their interaction, as well as coded variables for the music conditions (preferred vs unpreferred [1, 0] and familiar vs unfamiliar [1, 0]). Participant ID was included as random variable. The full formula was: *power ∼ enjoyment rating * familiarity rating + preference category + familiarity category + (1 | Participant ID)*. This resulted in coefficient maps for all predictors, channels and frequencies. Coefficients were then corrected for multiple comparisons using a cluster-based permutation approach. A null-distribution of 1,000 random permutations was created by randomly shuffling condition labels and repeating the linear mixed-effects modelling (note that only 1,000 permutations were considered computationally feasible, given that the analysis encompassed 64 channels x 119 frequencies). As in the condition comparisons above, alpha thresholds were again *p* = .05. The minimum number of adjacent electrodes was again three and the minimum number of frequency bins was five.

For MI, a linear mixed-effects model was fitted for each channel, separately for the delta (0.25–4 Hz) and theta (4–8 Hz) band. Outcome variable was the averaged MI. The same model formula and parameters were used as for spectral power permutation analyses. To give an indication of effect sizes for mixed model results, the peak beta coefficient for each cluster is reported.

## Results

### Behavioural results

As expected, songs in both the all-time and recent favourite conditions received high enjoyment ratings (see Figure 1; all-time favourite: *M* = 94.0, *SD* = 8.1; recent favourite: *M* = 93.3, *SD* = 7.6), with no significant difference between them (*t*(31) = 0.55, Cohen’s *d =* .097, *p*_FDR_ = .70). In contrast, tempo-matched songs from disliked genres showed greater variability and were rated substantially lower in enjoyment (all-time-matched disliked: *M* = 40.6, *SD* = 22.8; recent-matched disliked: *M* = 41.8, *SD* = 26.7), but again with no significant difference between them (*t*(31) = -0.19, Cohen’s *d = -*.03, *p*_FDR_ = .85). Importantly, the favourite conditions were rated as significantly more enjoyable than the disliked conditions (all Cohen’s *d* > 1.84, all *p*_FDR_ < .001).

**Figure 1.**
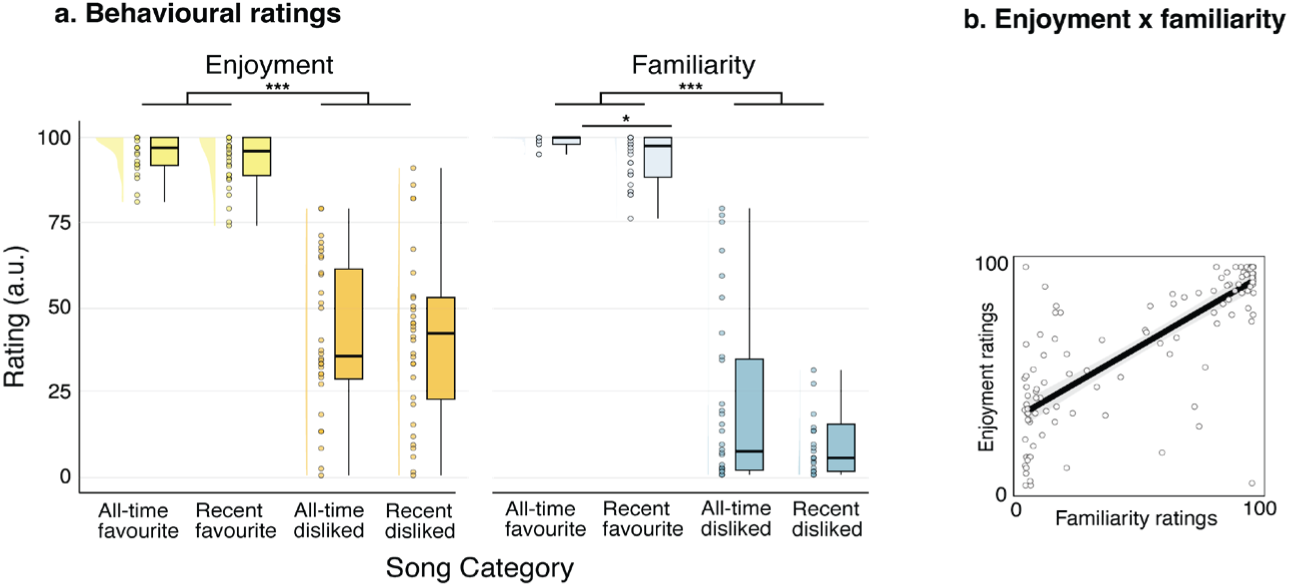
A) Behavioural ratings of enjoyment and familiarity across song categories. Distribution of participant ratings for the four song categories: all-time favourite, recent favourite, all-time disliked, and recent disliked. Each plot combines a density distribution (half-violin) on the left, jittered individual data points in the middle, and boxplots indicating the median and interquartile range on the right. **Enjoyment (Left Panel):** There was a highly significant main effect of preference (*p* <.001), where both "favourite" categories received near-ceiling enjoyment ratings compared to the "disliked" categories. No significant difference in enjoyment was observed between all-time and recent favourites. **Familiarity (Right Panel):** Favourite songs were significantly more familiar than disliked songs (*p* <.001). Notably, a significant difference was found between the two favourite categories (*p* <.05), with all-time favourite songs being rated as more familiar and showing a tighter distribution at the ceiling compared to recent favourite songs. *** denotes *p* <.001; * denotes *p* <.05; a.u. = arbitrary units. **B) Relationship between enjoyment and familiarity.** A linear mixed-effects model was used to test the association between enjoyment and familiarity ratings. Familiarity predicted enjoyment: the more familiar a song was, the more it was enjoyed.

Familiarity ratings followed a similar pattern. Both all-time and recent favourite songs received high familiarity ratings (all-time favourite: *M* = 96.8, *SD* = 8.6; recent favourite: *M* = 92.6, *SD* = 9.9); however, all-time favourites were rated as significantly more familiar than recent favourites (*t*(31) = 2.9, Cohen’s *d =* .51*, p*_FDR_ = .013). Tempo-matched disliked songs showed substantial inter-individual variability and were rated considerably lower in familiarity (all-time-matched disliked: *M* = 20.3, *SD* = 26.1; recent-matched disliked: *M* = 16.6, *SD* = 25.3, no significant difference, *t*(31) = 0.55, Cohen’s *d =* .098, *p*_FDR_ = .874). As with enjoyment, the favourite conditions were significantly more familiar than the disliked conditions (all Cohen’s *d* > 2.58, all *p*_FDR_ < .001).

We also investigated the relationship between familiarity and enjoyment ratings, using a linear-mixed effects model with enjoyment as outcome, familiarity as fixed effect, and random intercept for participant. This showed that familiarity ratings significantly predicted enjoyment ratings (*β* = 0.828, *SE* = 0.048, *t*(124) = 17.24, *p* < .001). Compared with a random-intercept-only model, including familiarity ratings significantly improved model fit (likelihood-ratio test: *χ*²(1) = 156.70, *p* < .001). This means that the more familiar a song was, the more it was enjoyed.

### Comparisons of spectral power between music conditions

An initial comparison of spectral power across all four conditions revealed differences between the all-time favourite and recent favourite conditions. Cluster-based permutation testing identified one significant cluster. This negative cluster emerged across widespread electrode sites in the low-frequency range spanning 3.00–5.25 Hz (*p*_Cluster_ = .006; Cohen’s *d*_peak_ = 1.47). The cluster showed higher delta-theta power during listening to recent favourite compared to all-time favourite songs. No other comparisons yielded significant clusters.

### Enjoyment and familiarity predict spectral power in distinct frequency bands

Next, we performed linear mixed-effects model analyses for all channels and frequencies to assess whether enjoyment and familiarity ratings and their interaction predicted spectral power (Figure 3). A cluster-based permutation approach was then used to correct for multiple tests across channels and frequencies. The main effect of enjoyment yielded five significant spatio-spectral clusters. Three were found in the theta range, and two in the beta range. The clusters in the theta range were found between 6 and 7 Hz, (*p*_Cluster_ < .001; *β*_peak_ = 0.76; 14 right-posterior electrodes), between 6.25 and 7.25 Hz (*p*_Cluster_ < .001; *β*_peak_ = 0.69; 10 frontal electrodes), and between 7 and 8 Hz (*p*_Cluster_ < .001; *β*_peak_ = 0.45; 6 left-temporal electrodes). The clusters in the beta band were found in the low beta range in posterior channels, between 13.75 and 15 Hz (*p*_Cluster_ *<* .001; *β*_peak_ = 0.67; 15 frontal electrodes), and in the higher beta range between 23.50–26.50 Hz (*p*_Cluster_ < .001; *β*_peak_ = 0.73; 14 posterior electrodes). All effects were positive, suggesting that power in the theta and low/high beta bands increased with more enjoyment.

The main effect of familiarity yielded one significant spatio-spectral cluster in the alpha range. This left-frontal cluster was found between 9.25 and 10.25 Hz (*p*_Cluster_ < .001; *β*_peak_ = 0.66; 14 electrodes). This positive cluster suggests that alpha power increased with more familiarity. The interaction between enjoyment and familiarity yielded two clusters, both in the theta range. The first one covered frequencies between 6 and 7.25 Hz (*p*_Cluster_ < .001; *β*_peak_ = 0.46; 10 central electrodes) and the second one covered frequencies between 6.5 and 7.5 Hz (*p*_Cluster_ < .001; *β*_peak_ = 0.55; 6 posterior electrodes). Both clusters suggest that for high-familiar songs, theta power increased with enjoyment (as seen in the main effect of theta above) but this effect was absent for low-familiar songs.

### Cortical tracking results

We first compared cortical tracking between all conditions (separately in delta and theta bands) using dependent *t*-tests with cluster-based permutation to account for multiple comparisons. Although music from the all-time disliked condition seemed to be tracked the most (see supplemental Figure S1), there were no significant clusters indicating systematic tracking differences between any of the conditions in the delta and theta bands. For an overview of MI values for each participant and across all frequencies, see supplemental Figure S2. Next, we also tested the relationship between participants’ ratings and cortical music tracking, equivalent to the power analyses above. For this, we used the previously described linear mixed-effects models to understand if and how enjoyment and familiarity predicted music tracking. This was done separately for tracking averaged within the delta and theta frequency ranges. We found that cortical delta music tracking was predicted by participants’ enjoyment ratings (Figure 4A) in one cluster (*p*_Cluster_<. 001; *β*_peak_ = −0.75; 3 electrodes). This negative cluster was located in right parieto-occipital channels and indicated that MI was stronger with less enjoyment. We did not find a main effect of familiarity ratings, enjoyment category or familiarity category on delta music tracking. However, the interaction between enjoyment and familiarity on music tracking was significant, with a negative cluster in fronto-central electrodes (*p*_Cluster_ < .001; *β*_peak_ = −0.52; 3 electrodes). Specifically, cortical tracking decreased with enjoyment for more familiar songs and increased with enjoyment for more unfamiliar songs.

For theta music tracking (Figure 4A), categorical enjoyment predicted tracking in a positive central cluster (*p*_Cluster_<. 01; *β*_peak_ = 1.68; 3 electrodes), suggesting that theta tracking was higher in the preferred-song conditions. Similarly, categorical familiarity predicted tracking in a negative central cluster (*p*_Cluster_<. 01; *β*_peak_ = -0.14; 5 electrodes), suggesting that theta tracking was higher in the less familiar conditions. However, neither the main effects of continuous enjoyment and familiarity was significant, nor their interaction.

## Discussion

The current study aimed to study the effects of familiarity and enjoyment on music processing when listening to preferred self-selected music and tempo-matched music from disliked genres. Including both all-time and recent favourite songs allowed us to assess the effects of familiarity and enjoyment in a systematic manner.

We hypothesised that listening to favourite songs would result in higher power in the theta, alpha, beta and gamma band compared to disliked songs, however, we did not find any differences when comparing all-time and recent favourite songs to their disliked counterparts. Instead, condition comparisons for spectral power revealed that listening to all-time favourite songs was associated with weaker delta/theta power than listening to recent favourite songs, potentially relating to familiarity. The tempo-matched song categories (‘all-time disliked’ and ‘recent disliked’ conditions) showed expected variability in familiarity and enjoyment, which likely is the reason why comparisons across conditions did not yield more significant results. Therefore, a mixed model approach was adopted using individual ratings for familiarity and enjoyment for all songs. These results did not confirm our finding in the delta/theta band. However, they revealed an effect of familiarity in the alpha band; the more familiar the song, the stronger was alpha power. Additionally, our mixed models revealed an effect of enjoyment on spectral power in the beta and theta bands. The more enjoyed a song was, the higher the beta and theta power. An interaction effect between enjoyment and familiarity on theta power revealed that the main effect of enjoyment was evident for high-familiar songs, but not for low-familiar songs.

We also found stronger cortical music tracking in the delta band when songs were less enjoyed, with no effect of familiarity on its own. However, we did observe an interaction between enjoyment and familiarity, showing strongest tracking when a song was familiar but not enjoyed. There were no significant differences in music tracking when comparing across conditions.

Further, a linear mixed-effects model revealed that familiarity predicted enjoyment of songs. This is in line with the well-established phenomenon of liking familiar music, sometimes referred to as the ‘mere exposure effect’ (Peretz et al., 1998; Madison & Schiölde, 2017).

### Low-frequency power distinguishes between recent and all-time favourite songs

We found a condition difference in delta-theta power between recent favourite and all-time favourite songs, with stronger low-frequency power during the recent favourite song than the all-time favourite song (Figure 2). This might suggest that low-frequency power could be associated with familiarity, however our mixed model results using continuous ratings did not replicate this finding. We are therefore careful to draw any firm conclusions on this result in low frequency bands.

**Figure 2.**
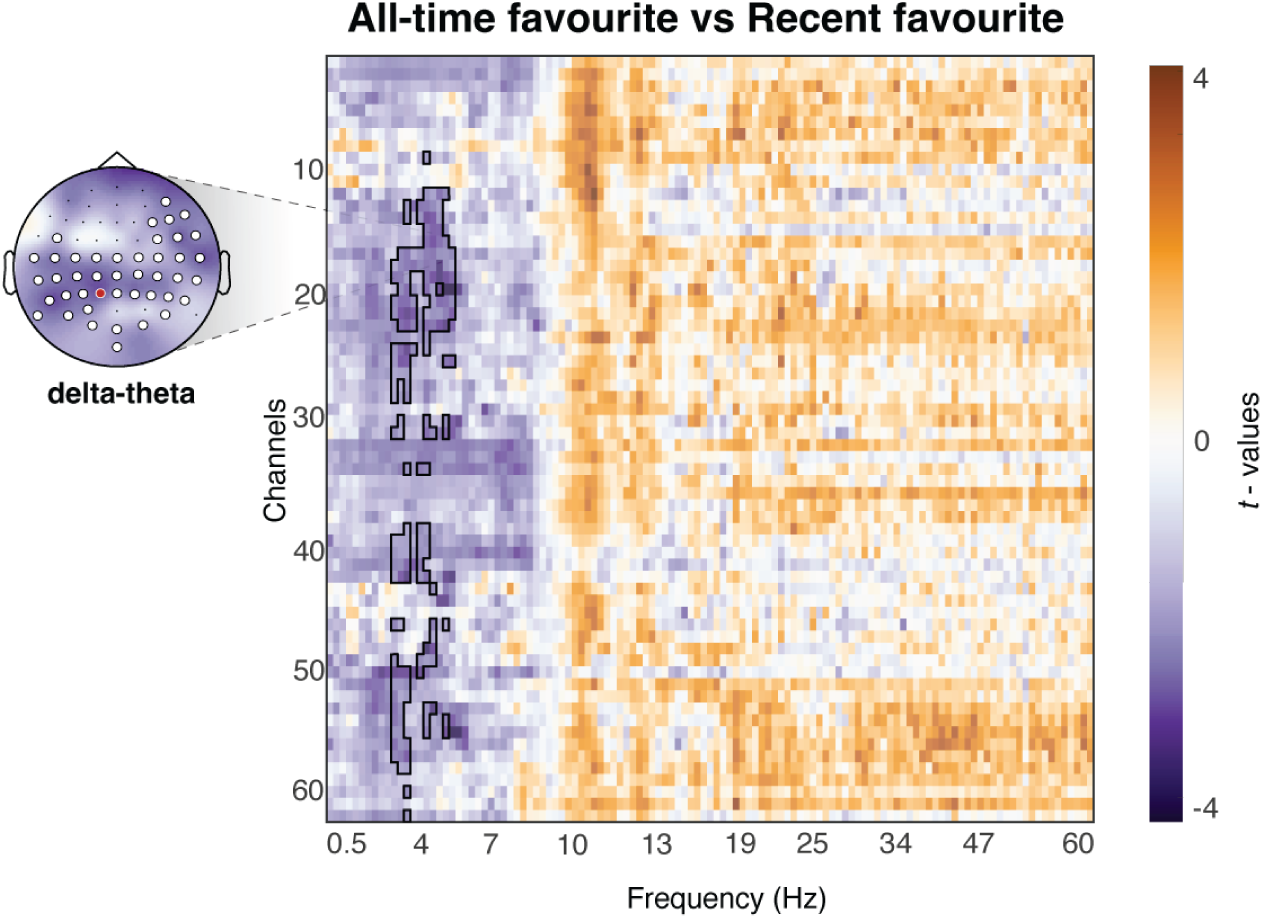
Spectral power comparison between all-time favourite and recent favourite music conditions. A channel-by-frequency map of t-values resulting from dependent *t-*tests. One significant cluster (outlined in black) was identified using cluster-based permutation. The negative cluster was observed in the low-frequency range, within the delta-theta band (3.00–5.25 Hz), indicating significantly higher spectral power for recent favourite songs compared to all-time favourite songs across widespread electrodes. The topographical map displays the spatial distribution of the averaged *t*-values within each significant cluster (white circles indicate significant electrodes with the peak effect indicated in red with white outline).

There is some evidence that supports this finding. Stronger theta power to unfamiliar than familiar music has been found (Thammasan et al., 2016; Meltzer et al., 2015). Theta oscillations have been shown to be involved in the encoding of new information (Klimesch, 1999), increased working memory demands (Klimesch, 2012), error processing as well as predictive processes (Arnal & Giraud, 2012; Kučikienė & Praninskienė, 2018; Recasens et al., 2018). Thammasan et al. (2016) reasoned that higher theta power during unfamiliar songs is due to participants paying more attention and to reflect working memory and episodic memory processes. Our results seem to support this. However, it is important to note that while stronger theta power seemed to be connected to the less familiar songs, both favourite songs in our study were still familiar, but one was more familiar than the other (Fig. 1). Literature on familiarity and delta power is less clear. One study suggested stronger delta power when listening to familiar compared to unfamiliar songs (Thammasan et al., 2016). However, they did not account for enjoyment. It is possible that our results indicate a specific effect of familiarity for highly enjoyed songs, but as stated before, we cannot draw any firm conclusions as this effect was not replicated in our mixed-model analyses.

### Alpha power indexes familiarity

Our results indicated that listening to familiar music is associated with an increase of alpha power. We found this effect as an effect of self-rated familiarity, when controlling for enjoyment. This finding is in line with Thammasan et al. (2016), who reported stronger alpha power when participants listened to 2-minute excerpts of familiar compared to unfamiliar songs. However, the opposite pattern has also previously been found, with familiar music associated with reduced alpha power (or increased alpha desynchronisation) (Malekmohammadi et al., 2023). A key methodological difference was that this study used short (10-second) excerpts, whereas our study, as well as Thammasan et al. (2016), employed longer musical pieces (2 minutes). It is therefore possible that the direction of alpha effects depends on the time scale, with sustained listening to familiar music being associated with increased alpha power.

We initially expected our results to be an indication of predictive processes. However, previous literature shows that predictive processes are related to a reduction in alpha power instead, specifically in the context of predictive sounds (Mohanta et al., 2021). Increased alpha power during music listening (relative to rest) has been associated with internally directed attention, such as mind-wandering, imagination, and meditative states (Markovic et al., 2017). Extending this interpretation to our findings, it is possible that greater familiarity facilitates this inward shift, leading to increased alpha power. In line with this, a prominent account of alpha oscillations proposed that alpha synchronization reflects the active inhibition of task-irrelevant processing and the controlled access to stored information (Klimesch, 2012). Under this framework, increased alpha power during more familiar music listening may indicate greater reliance on internally generated representations and memory-related processes supported by long-term knowledge of the musical material (Peretz et al., 1998).

### Theta and beta power index enjoyment

We found an effect of enjoyment on theta and beta power, with more enjoyed music associated with increased theta and beta power. An association between theta power and music evoked pleasantness is well established (e.g., Ara & Marco-Pallarés, 2020, 2021; Sammler et al, 2007; Ara et al., 2026).

Frontal theta power has consistently been implicated in emotional processing. For example, it has been shown that listening to a participant’s favourite song elicited higher theta, alpha, and beta power, while no differences emerged between neutral and moderately preferred music (Tseng, 2021). This suggests that strong subjective preference may be necessary to elicit pronounced prefrontal theta responses. Another study found increased fronto-central theta activity during listening to pleasurable music, with theta power scaling alongside arousal and emotional intensity (Chabin et al., 2020). These findings have been interpreted as a role of frontal theta power in reward processing and affective engagement with music. A limitation of the current study is that participants’ emotional responses were not measured, making it unclear whether the observed theta power increase indeed reflects emotional processing.

We also found an interaction effect of enjoyment and familiarity on theta power. Specifically, the theta increase associated with higher enjoyment seemed to have been driven by songs with high familiarity (Figure 3). This interaction, and the fact that in the condition comparisons the only differences found were between the two favourite song categories, could indicate that enjoyment effects on theta power are strongest for familiar music. This suggests that when investigating the effect of enjoyment on music processing, it is essential to assess subjective familiarity with the music.

**Figure 3.**
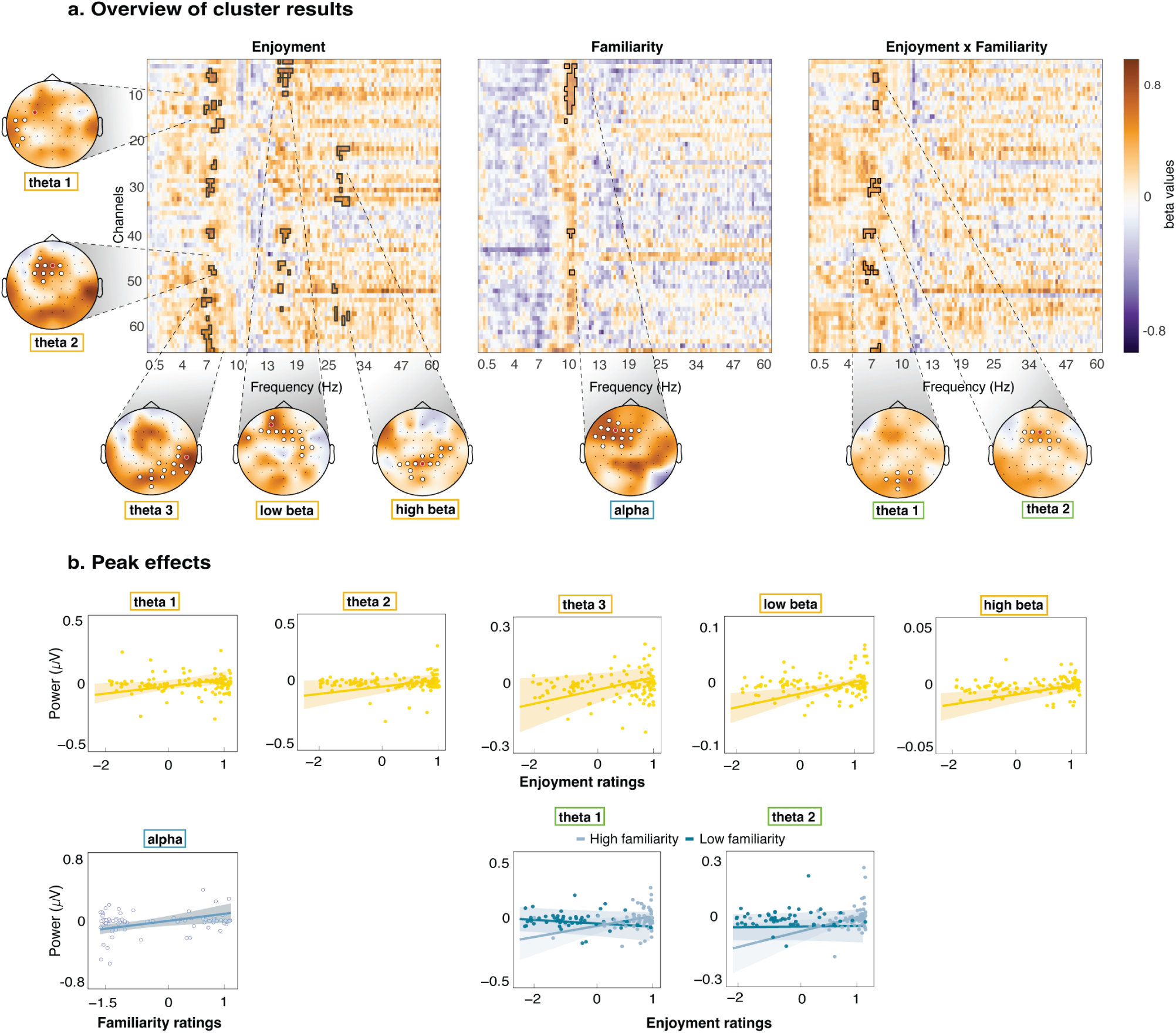
Effects of enjoyment and familiarity on spectral power. **a. Frequency and spatial distribution of spectral effects.** Channel-by-frequency *t*-maps with significant clusters outlined in black, showing how spectral power was modulated by enjoyment (left), familiarity (middle), and the enjoyment × familiarity interaction (right). Topographical maps show the spatial distribution of these *t-*values, with significant channels marked by white circles and the peak channel indicated in red. For enjoyment, positive clusters emerged in theta frequency bands with a frontal-midline distribution and a right-posterior distribution, as well as in low and high beta bands with central and right-parietal distributions respectively. These indicate that power increased with more enjoyment. For familiarity, a left-frontal cluster was observed in the alpha range, suggesting that alpha power increased with familiarity. For the enjoyment × familiarity interaction, two theta clusters (one central, one posterior) were identified, suggesting that theta power increased with enjoyment when songs were familiar, but this effect was absent for low-familiar songs. **b. Peak effects showing relationships between spectral power and subjective ratings.** Peak effects of each identified cluster (red electrodes in the topographies). Each plot displays the predicted power values (y-axis) as a function of the corresponding predictor (x-axis), with shaded areas representing 95% confidence intervals. Dots represent individual datapoints.

The literature on beta power and music enjoyment is mixed. Beta power is commonly associated with predictive processes during listening (e.g., Morillon & Baillet, 2017), but less commonly with enjoyment. The above study by Tseng (2023) is in line with our findings of increased beta power when participants listened to their favourite music. Another study (Hadjidimitriou & Hadjileontiadis, 2012) found stronger beta power when listening to disliked compared to liked music, a pattern that is opposite of what we found. However, they used short 15-sec unfamiliar music clips, while the current study employed longer, partly self-chosen and familiar music clips. Prior work has also linked beta activity to reward processing and predictive timing (Mas-Herrero et al. 2015, Kawasaki and Yamaguchi 2013, Weigmann, 2017). For instance, it was proposed that enjoyment arises partly from the brain’s ability to anticipate rhythmic structure, with oscillatory activity supporting temporal prediction and reward (Weigmann, 2017). However, as we found no effect of familiarity on beta power when controlling for enjoyment, we cannot conclude that the effect of enjoyment was linked to prediction processes. Our results show that, when measuring both individual familiarity and enjoyment and using diverse stimuli, beta power was primarily linked to enjoyment.

### Subjective enjoyment predicts cortical delta tracking

We found that more enjoyment was associated with less cortical tracking when controlling for the familiarity of music (Figure 4). Our results are in line with others that showed that more enjoyment was associated with lower neural tracking of acoustic temporal features (Hartung et al., 2025). Interestingly, they found that tracking of higher-level predictions (such as melody) was instead linked to higher enjoyment and familiarity. They suggested that when tracking of higher-level predictions was not possible, the tracking was focused on lower-level acoustics and was thus related to lower enjoyment. It could be argued that one would be less able to predict melody when one is unfamiliar with a song, and this would explain why the tracking was focused on lower-level features. However, our analyses revealed an interaction effect of enjoyment and familiarity, which showed that the main effect of enjoyment was present only in high-familiar songs (Figure 4). Songs rated as high in familiarity but low in enjoyment may represent music that participants are frequently exposed to without actively choosing to listen to them. Popular songs, for example, are often encountered in social settings, on the radio, in advertisements, films, or through social media, leading to high familiarity without necessarily eliciting enjoyment. In low-familiar songs, the opposite pattern was found, where more enjoyment was associated with slightly stronger tracking. This is in line with a previous study showing increased cortical tracking with more subjective enjoyment, which only used unfamiliar stimuli (Keitel et al., 2025). Another study found no effect of enjoyment, though they analysed tracking of spectral flux which could potentially explain the difference (Weineck et al., 2022).

**Figure 4.**
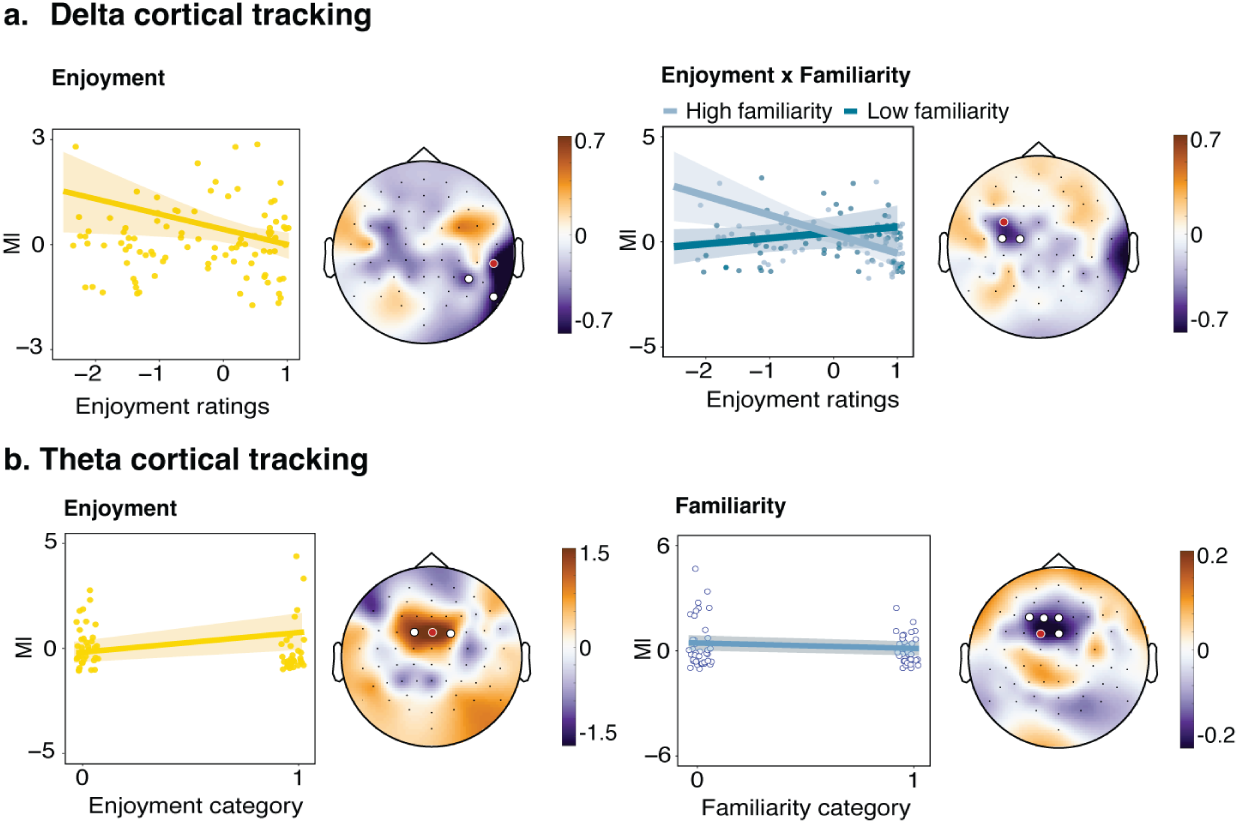
Linear Mixed-effects Model (LMM) results for cortical tracking. **a. Delta cortical tracking.** *Left panel – Main effect of enjoyment:* One parieto-occipital cluster found a negative relationship between continuous enjoyment ratings and delta cortical tracking (measured through Mutual Information; MI), where lower enjoyment ratings predicted higher tracking. *Right panel – Enjoyment × Familiarity interaction:* There was also a significant cluster for the enjoyment x familiarity interaction. For high-familiarity stimuli (light teal), there was a negative relationship between enjoyment and tracking, whereas for low-familiarity stimuli (dark teal), the relationship was reversed. The corresponding topographical map shows that this interaction effect was significant in a cluster of fronto-central electrodes (white circles). **b. Theta cortical tracking.** *Left panel – Main effect of enjoyment:* The scatter plot shows a positive relationship between enjoyment category (0 vs. 1) and theta MI, suggesting modulation of theta tracking by enjoyed versus non-enjoyed stimuli. The topographical map shows that the significant cluster (white circles) was located in a fronto-central region. *Right panel – Main effect of familiarity:* The scatter plot shows a negative relationship between familiarity category (0 vs. 1) and theta MI. The corresponding topographical map reveals a significant cluster of fronto-central electrodes (white circles). *Note*: Significant channels are marked by white circles, a red circle indicates the peak effect, for which the scatterplot is shown. The scatterplots display the predicted MI values (y-axis) as a function of the corresponding predictor (x-axis), with shaded areas representing 95% confidence intervals. Dots represent individual datapoints.

Previous studies have found stronger tracking of familiar music (Weickeck et al., 2022; Doelling & Poeppel, 2015) as well as stronger tracking of unfamiliar music (Kumangai et al., 2017, 2018) whereas we found no main effect of familiarity at all. Potentially the differences in stimuli and methods that were used to analyse tracking could be the reason for these contradictions. In Figure 4 (right panel) it can be observed that songs that were both highly familiar and highly enjoyed were tracked the least. One possible explanation is that these songs were more accurately predicted by listeners, reducing the reliance on acoustic information. Thus, resources are directed at more challenging processing steps, such as semantic processing (Arnal & Giraud, 2012). This interpretation is consistent with our finding that greater enjoyment was associated with reduced tracking. Attention and expectation are thought to modulate neural responses in opposite ways; where attention has been found to increase neural responses, expectation has been found to reduce them (Arnal & Giraud, 2012). However, as we did not find a direct relationship between familiarity and cortical tracking, we cannot draw direct conclusions about expectation effects. Our results show the importance of measuring both subjective enjoyment and stimulus familiarity when studying cortical tracking, as opposite effects can be found depending on the level of familiarity.

Both categorical predictors, enjoyment and familiarity, were significant predictors for cortical theta tracking. We refrain from interpreting these effects because they are not reflected in continuous ratings or condition comparisons. However, the present findings reveal a possible dissociation between delta and theta band tracking. Previous studies have often looked at delta and theta tracking jointly, as “low frequency tracking” (Doelling et al., 2019; Harding et al. 2019). Our results suggest that delta and theta tracking may reflect distinct processes and therefore should be analysed separately rather than being combined into a single low frequency tracking band.

### Limitations

Several limitations should be considered. Participants likely differed in how they defined and selected “all-time” and “recent” favourites, introducing variability in familiarity and emotional salience. The use of post-listening ratings may also have influenced neural responses, as rating tasks can alter EEG amplitudes across frequency bands (Markovic et al., 2017). Additionally, anticipation effects may have arisen for the preferred songs because participants submitted these songs themselves and could therefore anticipate hearing their preferred songs.

### Conclusions

The current study applies a novel paradigm, not just aiming to distinguish the neural correlates between liked versus disliked music but separating familiarity from enjoyment by using all time and recent favourite music. We highlight the importance of controlling for both enjoyment and familiarity to disentangle the different effects they have on neural activity, as this can lead to opposing results shown in the literature.

## Additional Information

### Author contributions

Conceptualization: S.M.V., A.K.; Formal analysis: S.M.V., R.H.T., T.A., A.K.; Data Collection: S.M.V., S.C.A., S.K., A.K., Writing - Original Draft: S.M.V., R.H.T, T.A., A.K., Writing - Review & Editing: S.M.V., R.H.T., T.A., S.C.A., S.K., A.K.; Visualization: T.A.; Supervision: A.K.; Funding acquisition: A.K.

## Data Availability

All data will be made available through the OSF (https://osf.io/9dkuh/files/).

## Supplementary Information

**Supplementary Table S1.**
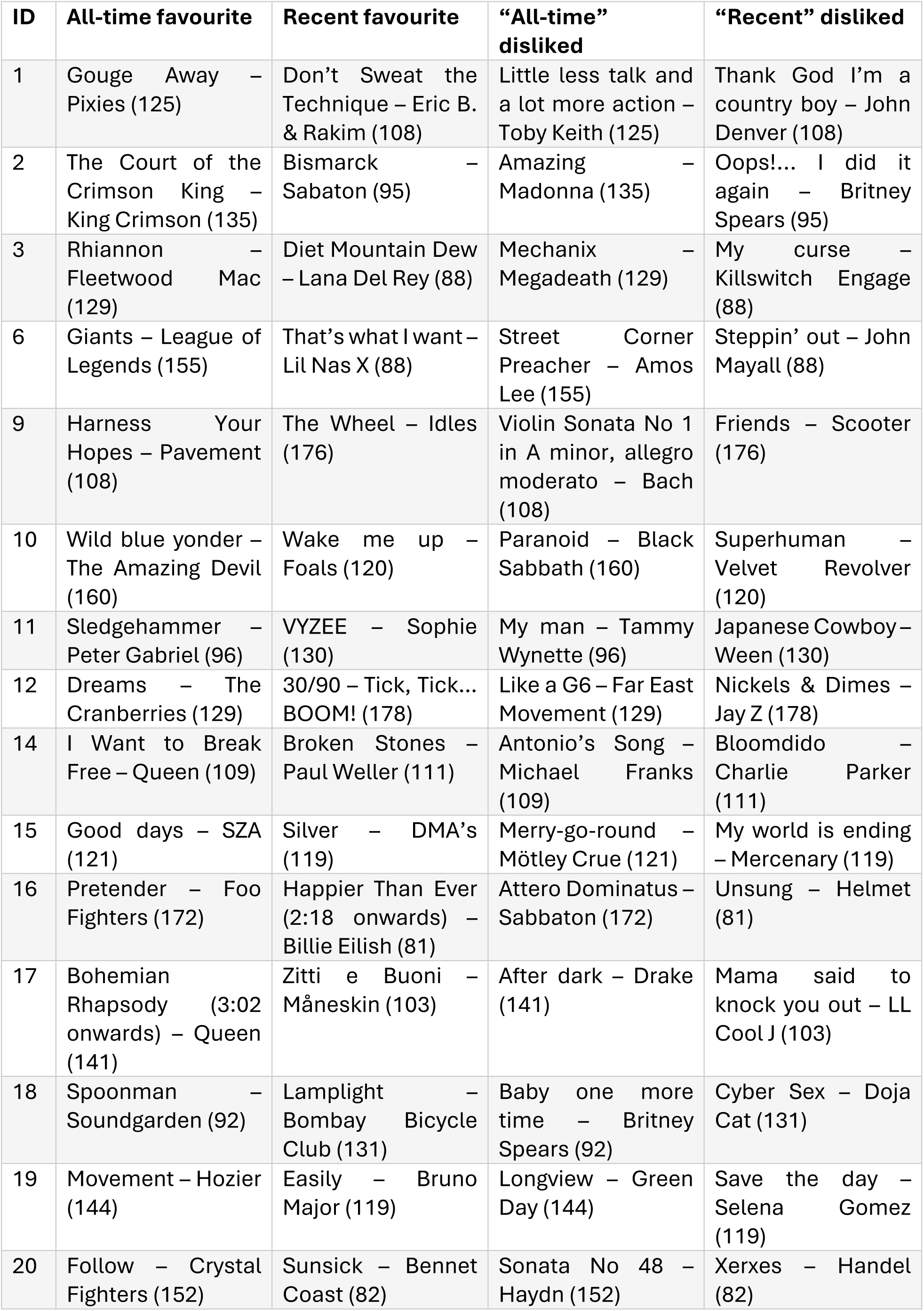
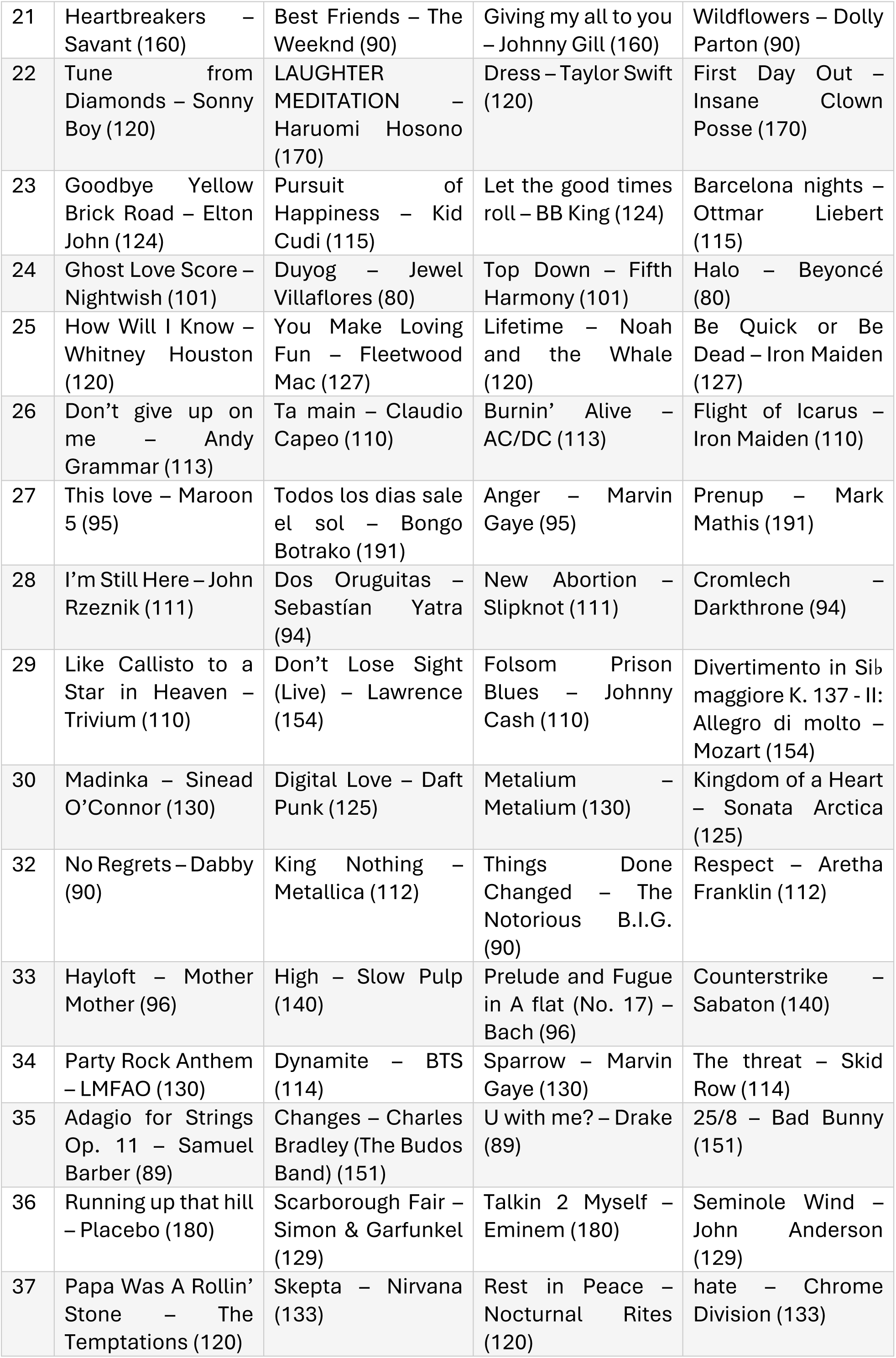
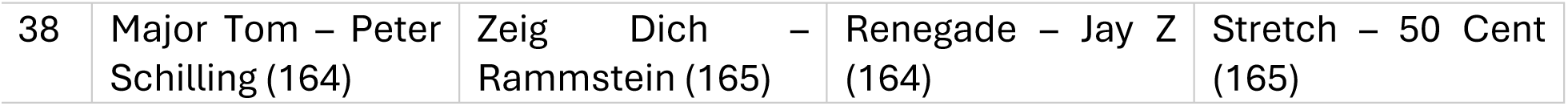
List of all songs. Tempo (in bpm) in brackets.

**Figure S1.**
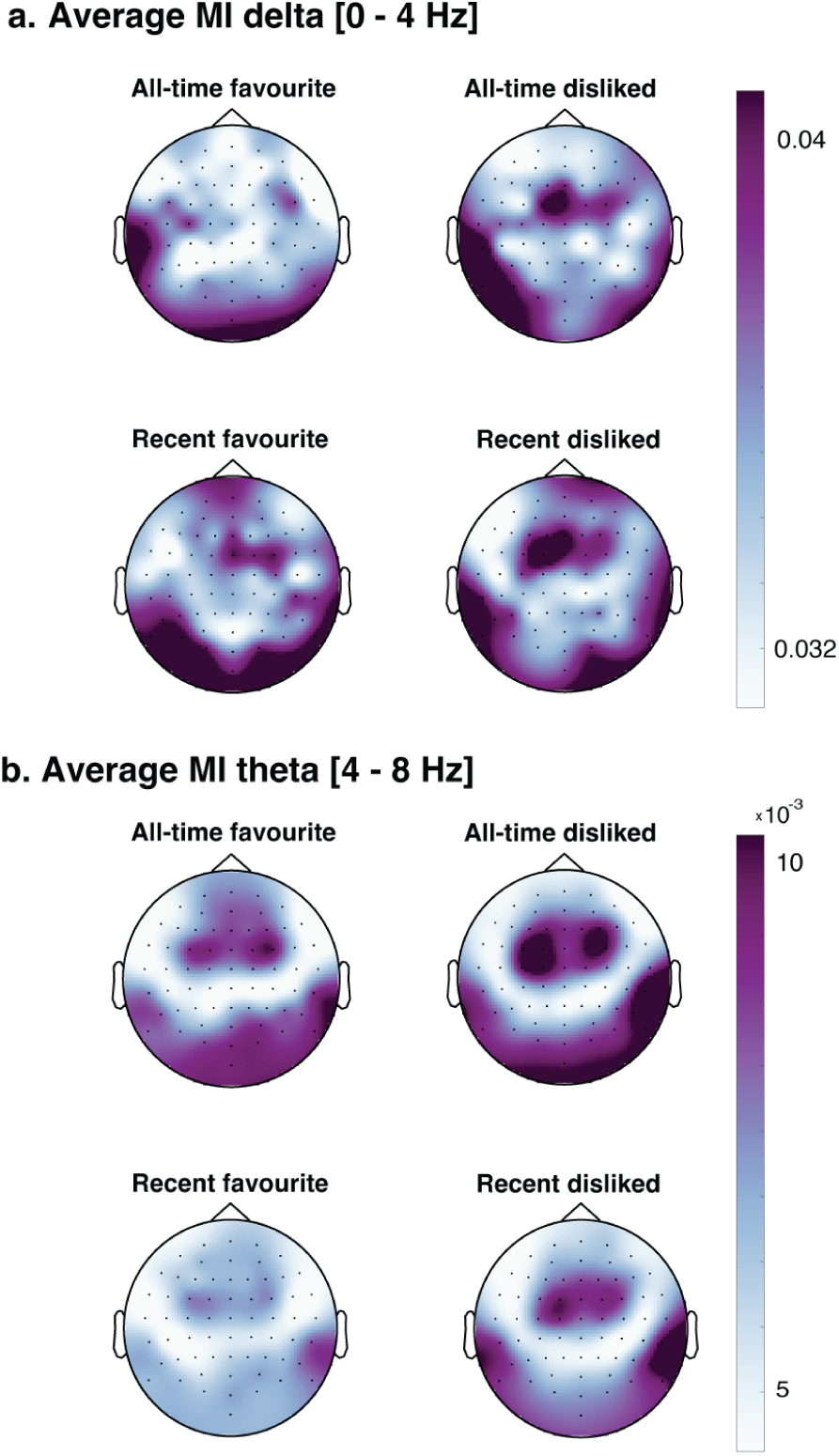
Topographical distribution of Mutual Information (MI). Scalp maps show the grand-average MI between EEG activity and the music envelope representation for the four experimental conditions (all-time favourite, all-time disliked, recent favourite, and recent disliked). MI was computed in the frequency domain (between 0.25 and 20 Hz, using a logarithmic frequency resolution) using complex wavelet coefficients of the music envelope and EEG signals. For each participant and condition, EEG data were aligned using the condition-specific delay that maximised MI, and MI was estimated between the complex EEG representation and a four-dimensional representation of the music envelope. Values were averaged across frequencies within the delta band (0–4 Hz; panel a) and theta band (4–8 Hz; panel b) and subsequently averaged across participants.

**Figure S2.**
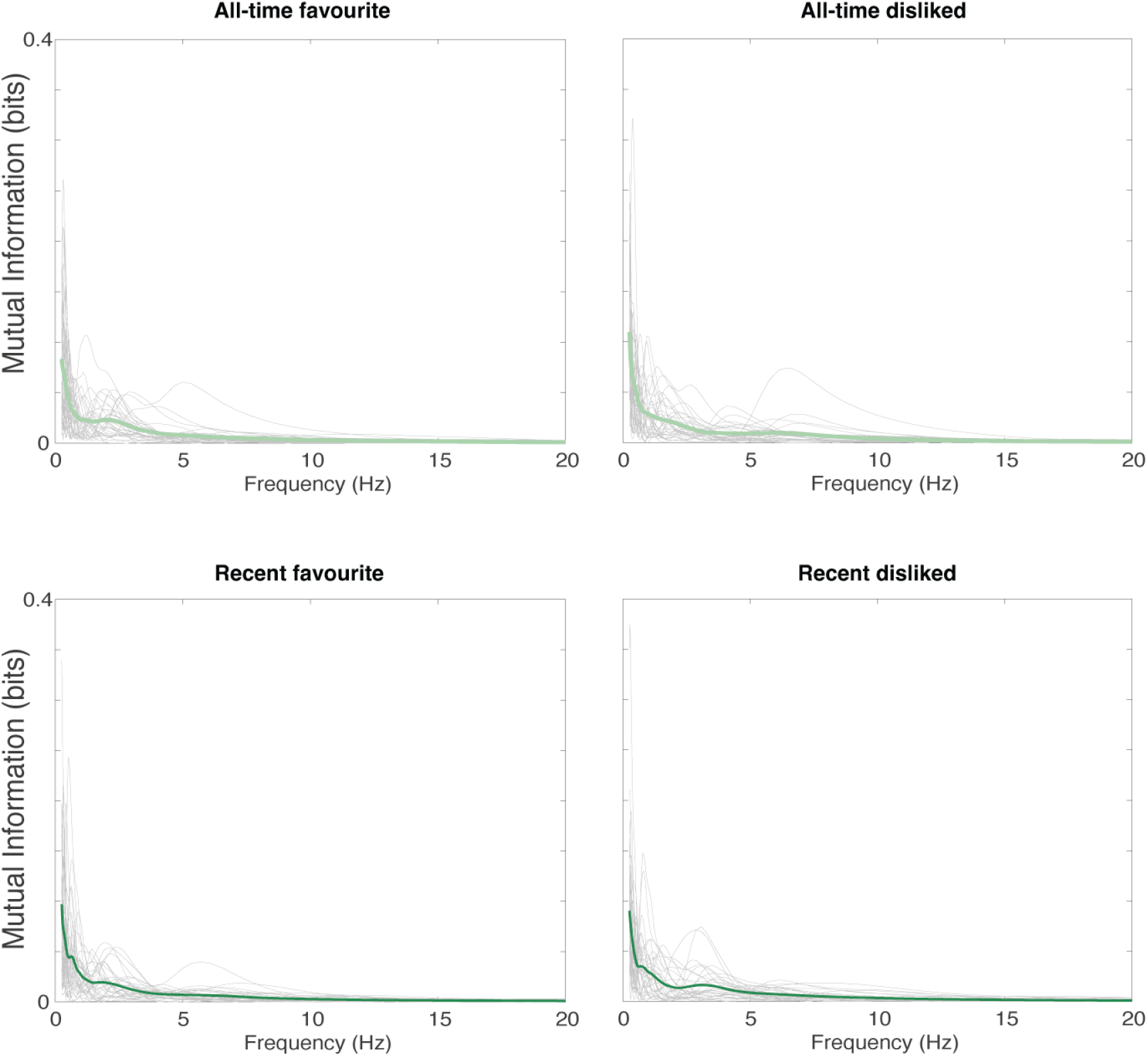
Frequency-resolved mutual information (MI) between EEG signals and music envelope for electrode Cz. For each listening condition (all-time favourite, all-time disliked, recent favourite, and recent disliked), MI was computed between the EEG signal and the music envelope representation in the frequency domain using the participant-specific optimal delay. Grey traces show individual participants’ MI spectra, while the coloured line represents the grand-average across participants. MI values are expressed in bits and plotted as a function of frequency (0–20 Hz). Across all conditions, MI was strongest at low frequencies, with a pronounced peak in the delta range (<4 Hz) and a rapid decline at higher frequencies. Above approximately 8–10 Hz, MI values approached zero for most participants, indicating that EEG–music coupling was primarily driven by slow temporal fluctuations in the music envelope.

